# Meta-analysis of Microbiome Association Networks Reveal Patterns of Dysbiosis in Diseased Microbiomes

**DOI:** 10.1101/2022.01.19.476958

**Authors:** Tony J. Lam, Yuzhen Ye

## Abstract

The human gut microbiome is composed of a diverse and dynamic population of microbial species which play key roles in modulating host health and physiology. While individual microbial species have been found to be associated with certain disease states, increasing evidence suggests that higher-order microbial interactions may have an equal or greater contribution to host fitness. To better understand microbial community dynamics, we utilize complex networks to study interactions through a meta-analysis of microbial association networks between healthy and disease gut microbiomes. Taking advantage of the large number of metagenomes derived from healthy individuals and patients with various diseases, together with recent advances in network inference that can deal with sparse compositional data, we inferred microbial association networks based on co-occurrence of gut microbial species and made the networks publicly available as a resource (github repository named GutNet). Through our meta-analysis of inferred networks, we were able to identify network-associated features that help stratify between healthy and disease states such as the differentiation of various bacterial phyla and enrichment of Proteobacteria interactions in diseased networks. Additionally, our findings show that the contributions of taxa in microbial associations are disproportionate to their abundances and that rarer taxa of microbial species play an integral part in shaping dynamics of microbial community interactions. Overall, this meta-analysis revealed valuable insights into microbial community dynamics between healthy and disease phenotypes.

## Introduction

The gut microbiome serves to provide a wide range of symbiotic functions, including metabolism, immune system development, and pathogen resistance [1]. While the gut microbiome plays an important role as a modulator of host health and disease, commensal colonizers are often susceptible to disruption, which has been shown to be associated with the development of disease states [2, 3, 4]. The advancement of sequencing technologies has fueled the rapid expansion of metagenomic data availability, enabling association studies between the human microbiome and various disease states [5, 6]. While many microbiome studies rely on differential analysis to identify individual bacteria of interest between cohorts, the ability of network analysis to provide high level insights into global and local structures makes it an attractive approach to study the dynamic nature of microbial communities.

Metagenomic co-occurrence has been widely applied in metagenomic studies to construct microbiome networks and better understand microbiome community structures [7, 8, 9]. However, features of metagenomic data pose several challenges to microbial co-occurrence analysis. Firstly, as sequencing technologies are not able to capture the true absolute microbiome abundance of samples, sequence abundances need to be represented as a proportion, rendering species abundance compositional by nature [10]. However, relative abundances, a common measure used to represent microbial abundances, is often considered a flawed metric to use in co-occurrence-based approaches due to the constant sum constraint, where assumptions of correlation metrics such as the independence between features are violated [11, 12]. As relative abundances of species are dependent on the relative abundances of every other species present, abundance values of a given sample are no longer independent of each other when normalized to relative abundances. As such, alternative methods of normalization or transformation of raw abundance values remain necessary to compare species co-abundances across samples of varying sequencing depths. Additionally, the use of compositionally aware association measures and methods are needed to handle the compositionality of microbiome datasets [13, 14, 15]. Various methods have been proposed to address the challenges of analyzing compositional data, and these methods that have been reviewed in detail [10, 14, 16, 17, 18, 19]. Secondly, microbiome data is often subjected to issues of sparsity, where microbiome abundance matrices are zero-inflated due to heterogeneity within and between samples. Rare taxonomic species and/or insufficient sequencing depths contribute to the sparsity often seen in microbiome datasets [20, 21]. The sparsity found in metagenomic datasets introduces challenges to log-ratio based transformation techniques used to handle compositionality. Additionally, correlations of sparse datasets can lead to strong spurious correlations [15, 20]. Non-parametric and ranked-based correlation measures such as Spearman’s Rho are also susceptible to multi-way ties due to matrix sparsity and heavy-tailed distributions, and quickly deteriorate in presence of many zeros [12, 22, 23]. Thirdly, indirect correlations can often add noise to correlation-based interaction inference methods, where these indirect associations (e.g. spurious associations) can be driven by indirect species associations, batch effect, or environmental factors [9, 15, 24, 25, 26].

Despite the challenges of utilizing co-occurrence metrics on metagenomic datasets, a wide range of methods have been adopted, developed, and utilized to better understand microbial associations. In general, methods used to study microbial associations can be grouped into two categories: (1) traditional/classical correlation methods (e.g. Pearson, Spearman, Kendall’s Tau), and (2) compositionally-aware methods. While compositionally-aware methods vary in their algorithms, they all seek to mitigate the confounding factors imposed by the current limitations of compositionality found in microbiome datasets. Compositionally-aware methods can be further sub-categorized into correlation-based methods (e.g. SparCC [27], CoNet [28], CCLasso [29]) and conditional dependence methods (e.g. SPIEC-EASI [24], Flashweave [25]) which try to differentiate between direct and indirect conditional dependencies). While the review and benchmarking of available methods’ performance remains beyond the scope of this paper, the discussion surrounding the complexities of various microbial inference techniques have been reviewed at length [7, 11, 12, 16, 20, 30, 31, 32]. As studies have previously shown, the results of networks generated from microbial association inference are largely dependent on the method used to infer the microbial interactions [32, 11, 30, 12]. Methods of interaction inference vary largely between studies in terms of accuracy and precision, and no one existing tool is able to address all issues of biases or confounding factors [11, 20, 25, 30, 32]. As there remains a lack of a community consensus and gold standard to evaluate the performance of methods used to infer microbial co-occurrence networks, users are largely left to decide the method of inference.

Variation between datasets come not only from intra-sample heterogeneity, but also different preprocessing and post-processing methods used between studies. The lack of consensus in computational methods, including annotation, quantification, preprocessing, and association methods makes comparison of findings between studies difficult. Despite the significant progress in methods development for compositionality-robust association methods and known issues with traditional correlation-based methods, traditional correlation methods (e.g. Spearman) still remains the most widely used type of association metric. The slow adaptation of compositionally aware methods for metagenomic data remains multi-factor and can likely be attributed to the exponential increase in computational requirements of compositionally aware methods, as well as legacy effect where researchers adopt the methods used in previous studies. Here in this study, we utilize the availability of large healthy and disease gut metagenome studies to preform a meta-analysis using microbiome association networks by re-analyzing and standardizing the analysis approach. By constructing association networks, we hope to be able to better understand microbiome community associations and community assembly dynamics within and between healthy and diseased microbial communities in an effort to identify features to help stratify disease states and potential microbial risk factors.

## Materials and Methods

### Datasets and Preprocessing

A curated list of sample accession numbers from publicly available human gut metagenome datasets was gathered from Gupta et al. [33] to be used this study. Samples included in this analysis total 4347 samples from 78 different study accessions, with samples spanning 10 different phenotypes including: Healthy, advanced (colorectal) adenomas (AA), atherosclerotic cardiovascular disease (ACVD), colorectal cancer (CRC), Crohn’s disease (CD), obesity (OB), overweight (OW), rheumatoid arthritis (RA), Type-2 diabetes (T2D), ulcerative colitis (UC). Samples from the following phenotypes included in Gupta et al., impaired glucose tolerance (IGT), symptomatic atherosclerosis (SA), and underweight (UW) were excluded from downstream analysis due to low sample count. Samples were downloaded from NCBI Sequence Read Archive via SRA Toolkit’s fasterq-dump. A summary of samples used in this study can be found in Table 1. The works of Gupta et al. focused the meta-analysis of gut microbiome species to develop a health status index that utilizes species-level gut microbiome profiling to stratify between microbiome health states. While we utilized a similar dataset to Gupta et al.’s study, there are several notable differences between our analysis approach. Firstly, we selected a Kraken+Bracken approach for microbial quantification and taxonomic assignment due to its superior performance compared to marker gene based methods as highlighted in a recent benchmark of metagenomic classification tools, where marker gene based methods ranked among the lowest among assessed tools in terms of precision and recall for species classification and lowest proportion of abundance quantified at species-rank [34]. Additionally, our meta-analysis focuses on species co-occurrences and network-based approaches rather than focusing on the prevalence of species-level abundances between samples. Lastly, while maintaining similar study accessions, we insured that all run accessions downloaded focused on available paired-end reads with the largest available spots rather than utilizing a mix of single-ended and paired-ended reads.

**Table 1:** Summary of gut microbiome datasets used in downstream gut microbiome association network analysis.

| Phenotype | # of Studies | # of Samples |
| --- | --- | --- |
| Healthy | 29 | 2568 |
| Advanced (Colorectal) Adenoma (AA) | 2 | 82 |
| Atherosclerotic Cardiovascular Disease (ACVD) | 1 | 152 |
| Colorectal Cancer (CRC) | 4 | 254 |
| Crohn’s Disease (CD) | 4 | 107 |
| Obesity (OB) | 15 | 324 |
| Overweight (OW) | 16 | 232 |
| Rheumatoid Arthritis (RA) | 1 | 92 |
| Type 2 Diabetes (T2D) | 3 | 236 |
| Ulcerative Colitis (UC) | 3 | 96 |

Samples were processed to remove low quality reads and Illumina adapters using Trimmomatic (v0.39) [35] with parameters SLIDINGWINDOW:4:20 LEADING:20 TRAILING:20 MINLEN:60. Trimmed samples were then mapped to the human genome assembly GRCh38 (hg38) using bowtie2 (v2.4.4) [36] to remove possible human read contamination from the metagenome samples. All remaining unmapped metagenomic reads were kept for downstream analysis. Additionally, low read count samples that were less than 1M reads were discarded from this analysis to prevent inclusion of under-sampled genomes. Distribution of the filtered reads can be found on Supplementary Figure 1. Following filter and trimming of samples, a total of 4143 (from 32 studies) out of the original 4347 samples (from 34 studies) were retained for downstream analysis. A complete list of accessions used in this analysis can be found in the GutNet repository.

### Microbiome Taxonomic Assignment and Abundance Quantification

Taxonomic assignment and species abundance quantification were preformed using Kraken2 (v2.0.8) [37]. The pre-built developer maintained ‘Standard’ Kraken2 database (version k2_standard_20201202) built on December 2, 2020 was used for taxonomic references. The Kraken2 database was built using RefSeq reference genomes, including references from archaea, bacteria, viral, plasmid, human, and UniVec_Core databases. Only archaea and bacterial counts were retained for downstream analysis. Kraken2 prokaryotic taxonomic assignments and abundances were then re-estimated with Bracken (v2.6.2) [38] for specieslevel re-estimation of abundances. Samples were aggregated into their representative disease phenotype to construct species level read abundance matrices.

### Species Abundance Processing and Filtering of Sparse Taxa

One of the challenges in dealing with metagenomic data for co-occurrence inference is the sparsity of metagenomic data. This sparsity can be attributed to a multitude of factors (e.g. sequencing depth, sample heterogeneity) and can cause spurious correlations and false-positives in statistical methods [11, 13]. To address some of the issues caused by matrix sparsity, we follow the suggestion of Wiess et al. [11] to filter rare taxa from abundance matrices based on 50% sample prevalence. Species-level abundance matrices were filtered to remove low prevalence taxa below a 50% sample prevalence threshold.

### Microbiome Association

Using prevalence filtered bracken reads count abundance matrices, species-level associations were inferred for each disease abundance matrices respectively. SPIEC-EASI [24] was selected as the association method for microbial association inference due to the method accounting for compositionality of microbiome data and potential indirect associations. SPIEC-EASI was run using the ‘MB’ method, a neighborhood selection method developed by Meinshausen and Bühlmann [39] used to infer sparse inverse covariance matrices from a network. SPIEC-EASI has been found to preform well in comparison with other association methods, and thus selected to be used in this analysis [15, 13, 25].

### Microbiome Network Construction

Microbiome co-occurrence network were constructed from association values computed using SPIEC-EASI, where values were filtered with a 0.1 absolute association value threshold. Network vertices were defined as prokaryotic species for species-level networks; vertices and node are used synonymously throughout. An undirected edge was constructed between two vertices if a significant association between two given vertices was inferred. Edge weights range between [-1, 1], where positive edges represent a positive association and negative edges represent negative associations. It should be noted that edge weights of conditional dependence methods cannot be directly compared to correlation based metrics and are not directly proportional even though their values are assessed on the same scale (e.g. Pearson, Spearman, SparCC) [24, 25]. Networks were visualized through Gephi [40] using Force Atlas 2 layout. All singleton nodes without edges were removed from the network.

In addition, a consensus network was constructed to analyze Proteobacteria interactions among the disease networks. Given an edge, if any vertices within that edge had an annotated Genus as Proteobacteria, the edge was kept. Utilizing all remaining edges, a consensus network for each disease was built, where the edge weight was equivalent to the number of networks containing each respective edge.

### Community Module Detection

Many methods developed for community module detection in network systems are based off of undirected, unsigned, and positive networks. However, methods for signed module detection remain largely under-explored. In many cases, negative edges are simply discarded or ignored. However, as microbiome interactions are highly-dynamic and involve not only positive interactions, it is important to maintain the use of signed interactions when possible. To address this challenge, we utilized the Leiden algorithm [41], which attempts to extend on the works of the Louvain algorithm [42]. The Louvain algorithm can sometimes have badly connected communities, whereas the Leiden algorithm guarantees that communities are well connected and locally optimized. The Leiden algorithm consists of three steps, first it performs a local moving of nodes, second it refines partitions, and lastly the aggregation of the network based on the refined partitions. The Leiden algorithm takes advantage of local moving procedure and is able to split clusters rather than only merging them as in the Leiden algorithm. Additionally, the Leiden algorithm is able to handle negative edge weights.

### Module Resilience

We proposed a resilience score to approximate the tendency of modules of gut bacterial species detected from the healthy microbiome network to remain in the same community in the gut microbiome associated with different diseases. Given a module i found in healthy network containing *r_i_* species, for each diseased network our approach finds the module in the diseased network *j* that contains the most members of the *r_i_* species (denote as *d_j_*) (so 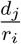 indicates the tendency of the species in module *i* staying in the same community in diseased network *j*). The resilience of module *i* is defined as the median of 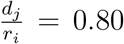, where *K* is the number of diseased networks (*K* = 9 in this paper). For example, module *i* contains 20 species, and 16 out of these 20 species are found in a module in the microbiome network for disease *j* (the remaining 4 species are found elsewhere), then *d_j_* = 16 and 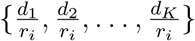. Assume 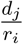 is 0.80, 0.90, 0.60, 0.70, 0.85, 0.75, 0.90, 0.35, 0.40 for *j* = 1,…,*K*, respectively, module *i* has a resilience score of 0.75 (the median). While this analysis was able to identify modules that were likely to be resilient to change, it does not provide information in regards how necessary the module was in regards to microbiome health nor does it identify ‘core’ microbiota, instead it shows how likely microbial species were to consistently form community modules across networks.

### Availability of the Programs and Inferred Networks

All network (GML) files, bioinformatics workflows and analysis scripts produced as part of this study can be found in a github repository named GutNet at https://github.com/mgtools/GutNet. Sample run accession numbers and associated study accession for all publicly available stool metagenome samples used in this study are available in the repository.

## Results

### Microbiome Composition and Sparsity Problem

The total number of species annotated in all datasets was 6463, spanning 4143 samples (see Table 1). When agglomerated at the Phylum level, we unsurprisingly found that Bacteroidetes, Firmicutes, Proteobacteria, Actinobacteria, and Verrucomicrobia were the 5 most dominant Phylum, with Bacteroidetes and Firmicutes dominating over 80% of the total relative abundance (Figure 1). This distribution of observed top Phyla is in line with previous studies that found similar distributions of top Phylum-level abundances in human gut microbiome [43, 44, 45].

**Figure 1:**
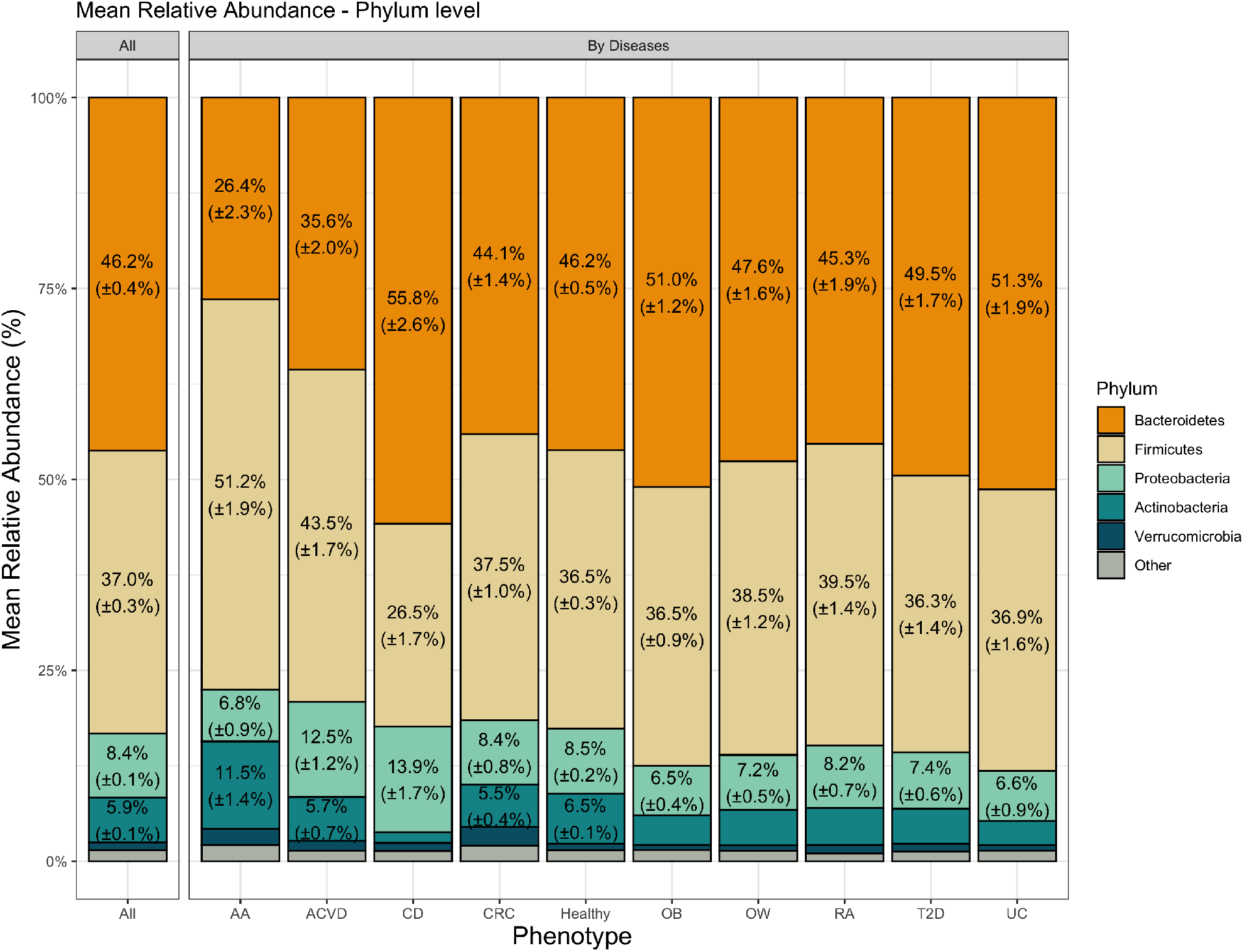
Mean distribution of species found within metagenome datasets by phenotype, agglomerated at the Phylum level. Numbers within each bar represent the mean relative abundance, accompanied by its standard error; only values above 5% are shown.

It has been shown that sparsity of microbial datasets affect correlation methods, and often result in spurious correlations. To address this issue, various explorations have proposed the use of filtering rare microbial taxa [11, 46]. Filtering species with low prevalence reduces the zero richness within datasets and helps resolve some of the statistical artifacts imposed by sparse datasets. Before deciding on a prevalence threshold, we evaluated the effect of imposing a prevalence threshold on microbial taxonomic distributions (Supplementary Figure 2). In most disease abundance matrices, the observed species present gradually decreases until approximately 65% prevalence, where thereafter the number of species post-filtering sharply decreases; the CD abundance matrix was the exception, where CD had a more linear relationship in terms of percent of species retained and percent of prevalence filtered.

We show that for all abundance matrices (except CD), a prevalence filter of 50% as suggested by [11] will result in a reduction in the number of species between 10.85% – 20.73% relative to the unthresholded datasets; with the exception of the CD dataset which will incur a 43.18% reduction in number of species observed after thresholding at 50% prevalence. Given the marginal differences in the number of species removed at prevalence values less than 50%, except for CD, we decided that a 50% species prevalence threshold was acceptable. Additionally, for the CD abundance matrix we decided that the trade-off of reducing sparsity was enough to warrant the loss of species present within the dataset, thus followed a 50% threshold for prevalence on all abundance datasets.

### Assessment and Comparison of Microbiome Ecological Diversity in Phenotype Specific Microbiomes

To evaluate the alpha-diversity between healthy and diseased microbiome datasets, we utilized the Shannon diversity index and species richness (observed number of different species) measures per each phenotype (Figure 2). For the alpha-diversity based on the species richness, we found that healthy datasets had a statistically significant different distribution compared to diseased datasets in terms of species richness (Figure 2A; two-sided Mann-Whitney U test, p-value= 2.9*e* – 4). Additionally, when testing the statistical difference between the healthy datasets versus each disease dataset individually, 8 out of 9 diseased phenotypes (i.e. AA, CD, CRC, OB, OW, RA, T2D, and UC) were found be statistically significant (Figure 2B; two-sided Mann-Whitney U test, p-value < 0.05). Shannon diversity measures between healthy and diseased datasets also showed a statistically significant different distribution (Figure 2C; two-sided Mann-Whitney U test, p-value = 1.6*e* – 12). Comparison between healthy and individual disease phenotypes also showed 5 out of 9 disease phenotypes (i.e. ACVD, AA, CD, T2D, UC) to be statistically significant (Figure 2D; two-sided Mann-Whitney U test, p-value < 0.05). Observations of significant differences in alpha diversity measures between healthy and diseased datasets are in line with previous studies that have used alpha-diversity measures as an indicator of disease-associated microbiome dysbiosis [47, 48].

**Figure 2:**
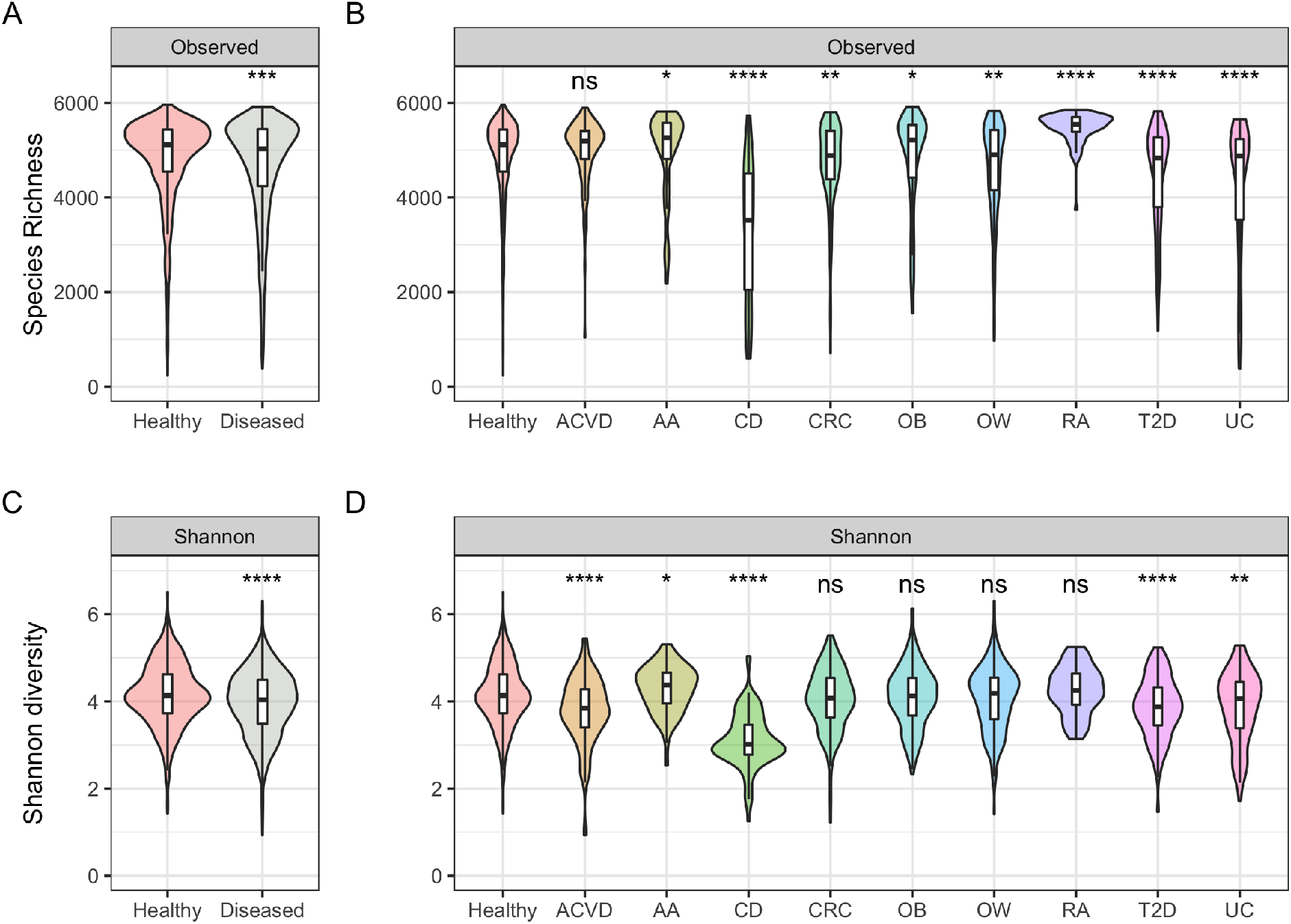
Alpha-diversity comparisons between datasets. (A) Species richness plot between healthy and diseased datasets, (B) species richness plot comparison between all phenotypes, (C) Shannon-diversity plot between healthy and diseased datasets, and (D) Shannon-diversity plot between all phenotypes. Two-sided Mann-Whitney U test was used to compare respective disease datasets against the healthy dataset. P-value significance are shown above violin plots; ns (non-significant; p-value > 0.05), * (p-value < 0.05), ** (p-value < 0.01), *** (p-value < 1*e* – 3), **** (p-value < 1*e* – 4).

For beta-diversity analysis, we used ordination plots to summarize the microbiome community data of healthy population and individuals with diseases. We used Bray-Curtis dissimilarity as the distance measure between the datasets, and used both t-SNE and NMDS approaches for dimensionality reduction. In the 2-dimensional ordination space shown in Supplementary Figure 3, samples with similar microbial compositions are close in the plots. The ordination plots show that samples did not cluster at the phenotypic-level, indicating that there is no discernible structure to microbiome abundance profiles that stratifies diseases purely based on taxonomic features.

### Microbial Association Network and Resilient Modules

To better understand microbiome associations and microbial community interactions in healthy and diseased gut microbiomes, we identified microbiome community modules within each microbiome network (Supplementary Figure 4-13). Co-occurrence networks were constructed for each phenotype, and community modules in each network were identified utilizing the Leiden algorithm. We compared the modules identified in the different microbiome networks to study the community module stability. By understanding the module resilience, we were able to identify microbiome community modules that were resilient to change, and identify species of bacteria that were more likely to be associated to each other regardless of the environment.

In our analysis, we were able to identify several modules of high module resilience. In many cases, modules of high resilience was populated by members of the microbiota in the same Clade. These include modules which were found to be Streptococcus-rich and Escherichia-rich at the Genus level, as well as Actinobacteria-rich and Proteobacteria-rich at the Phylum levels. Additionally, we also found modules with a mixture of Phyla that also exhibited high resilience, suggesting that resilience of modules may include both taxonomically assortative communities and those of mixed communities. While module resilience does not provide context as to why certain modules of microbial associations were retained through both healthy and diseased networks, it can help us better understand the underlying community structure and generate candidates for downstream hypothesis testing.

### Contributions of Taxa in Microbial Association Networks are Disproportional to their Abundances

By examining the species (i.e., the nodes) and their interactions (i.e., the edges) in the microbial association network, we can study their contribution to microbial community assembly. Analyzing the nodes of constructed association networks, we found that the top Phyla in each association networks comprised of Proteobacteria, Firmicutes, Actinobacteria, Bacteroidetes, Euryarchaeota, and Cyanobacteria (Supplementary Figure 14). However, the most abundant Phylum was that of Proteobacteria, which represent 42.37% of the total nodes found in the SPIEC-EASI association networks. This is in contrast to Bacteroidetes and Firmicutes which together only represented 26.89% of the total nodes found in SPIEC-EASI association networks. This contrast was surprising when compared to the distribution of microbial relative abundances, where Bacteroidetes and Firmicutes together represented > 80% of the mean total relative abundances (Figure 1).

Taking a closer look at microbial interactions of gut microbiomes between healthy and disease datasets, we analyzed the Phyla distribution of edge associations within each network. Similarly to network nodes, Phyla distribution of edges also did not show preference to Bacteroidetes and Firmicutes despite the dominant proportion of both Phyla in terms of relative abundances. As many of the interaction edges between microbial members lie between less populous Phyla, this highlights the importance of rarer species of the microbiome.

Differentiating between positive and negative edges in the network, we analyze the differences within and between the microbiome networks (Figure 4). Of notable observations, both Bacteroidetes-Bacteroidetes and Firmicutes-Firmicutes interactions were enriched in healthy populations compared to their diseased counterparts. While Firmicutes and Bacteroidetes did not exhibit drastic mean abundance differences in most disease datasets compared to the healthy dataset, the decrease in self-Phylum interactions may suggest that Firmicutes-Firmicutes and Bacteroidetes-Bacteroidetes interactions play an important role in maintaining gut homeostasis. Additionally, we found that Proteobacteria-Actinobacteria associations were enriched in disease networks compared to the healthy network and may be a signature of microbiome dysbiosis.

**Figure 3:**
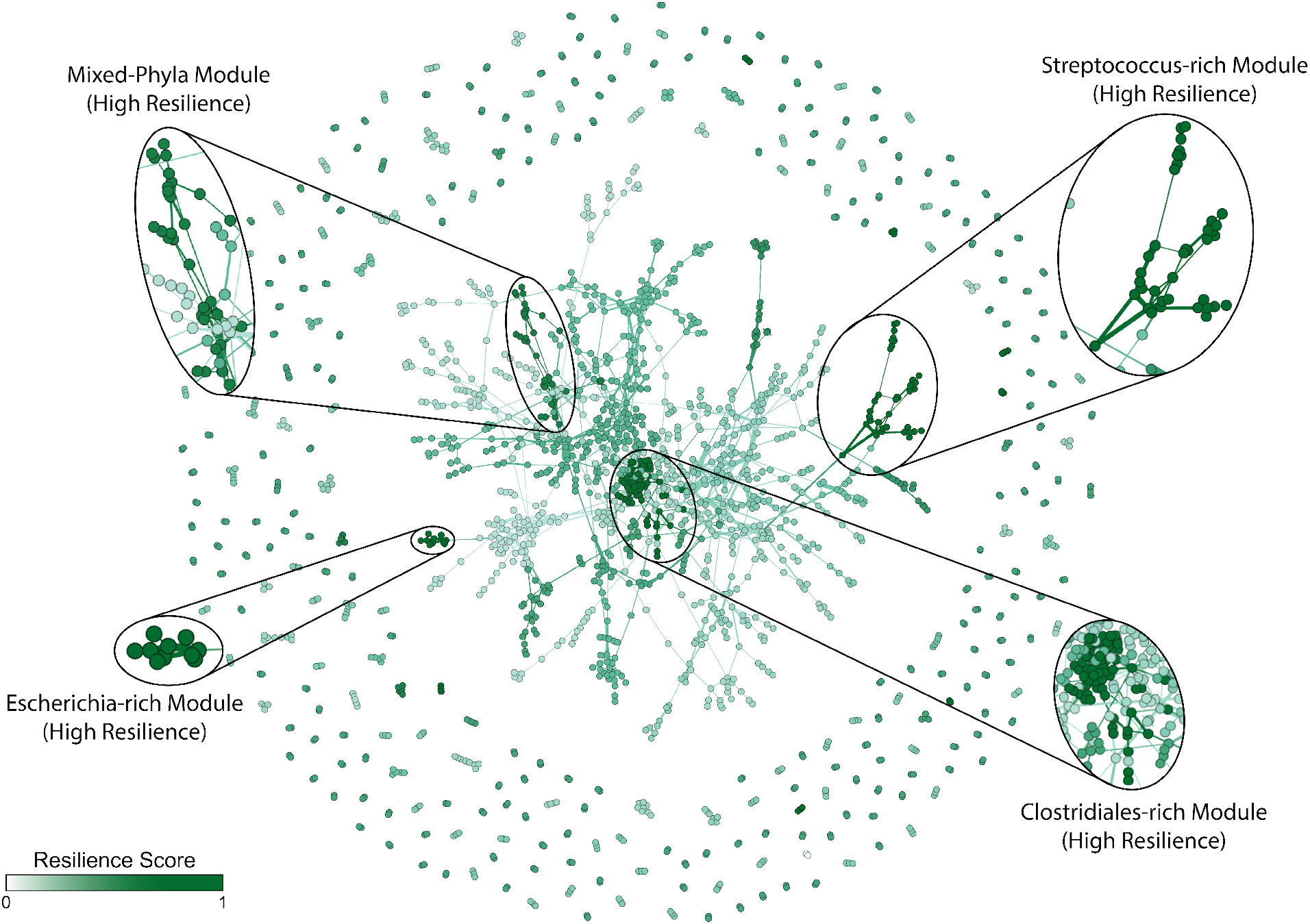
Microbiome association network, colored by module resilience. Module resilience scaled between [0,1] with lighter color modules represent lower module resilience, and darker color modules represent higher module resilience.

**Figure 4:**
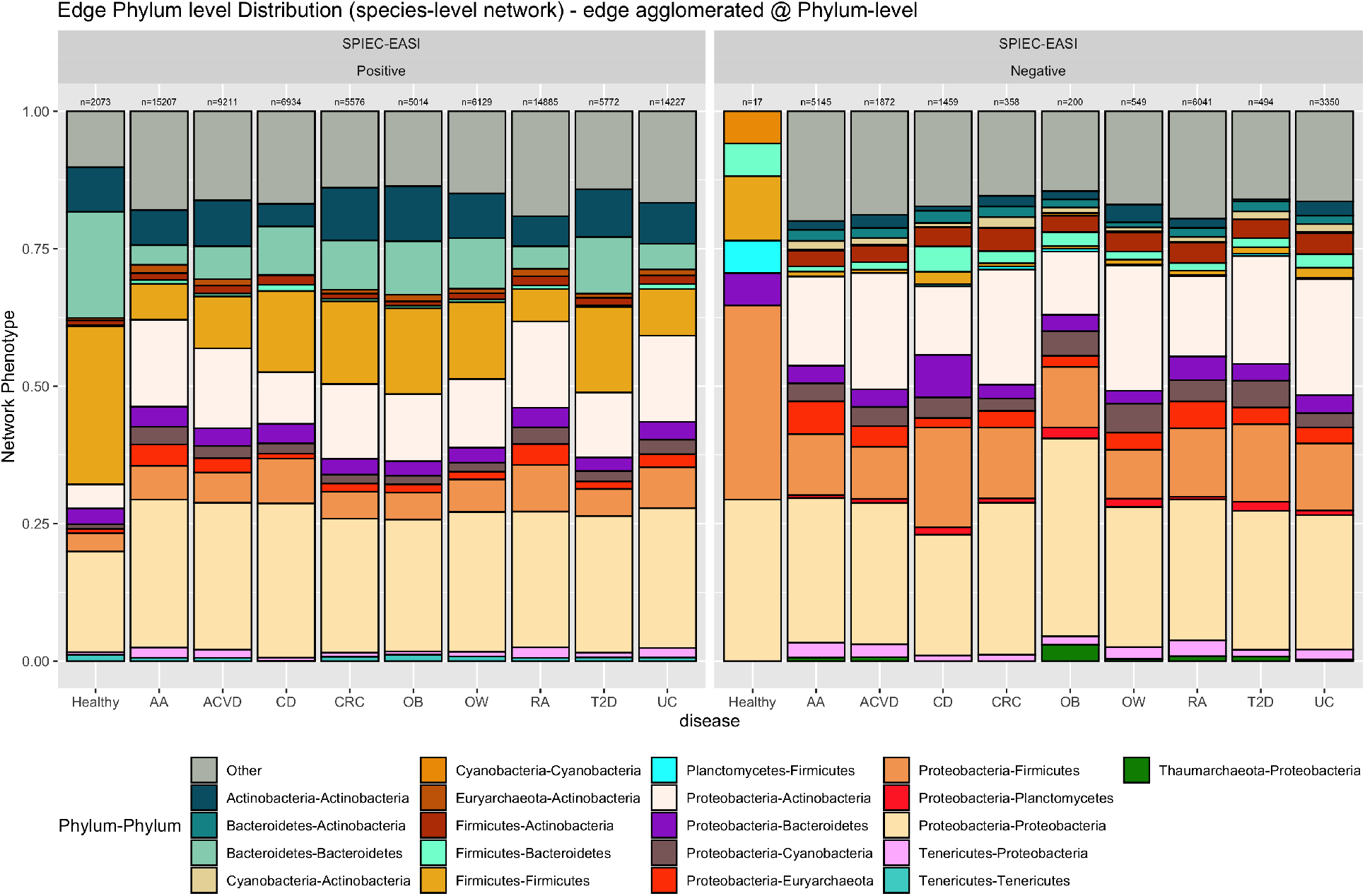
Taxonomic distribution (at the Phylum level) of the species involved in microbiome association networks by phenotype. (left) Positive edge distribution stacked barplots, (right) Negative edge distribution stacked barplots.

### Proteobacteria Interactions Enriched in Disease Association Networks

Previous studies have found that microbial abundances of Proteobacteria species were enriched in diseased microbiota and also have also proposed that Proteobacteria may be a signature of disease [49, 50]. While our results do not show consistent increase in mean relative abundance of Proteobacteria across all diseased datasets, only ACVD and CD datasets had a mean relative abundance greater than the healthy dataset (Figure 1), we observed that Proteobacteria participation in interactions (i.e., network edges) were significantly enriched in all disease networks. Proteobacteria was found to be the most dominant Phyla in terms of network edge participation, where Proteobacteria was part of either one or both vertices in a given network edge. On average, Proteobacteria participated in about 59% of the interactions in the microbiome association networks (healthy and diseased). Interestingly, the healthy network was identified as an outlier among networks (following Tukey’s method of outlier detection) with 33.88% of the interactions involving Proteobacteria (Figure 5).

**Figure 5:**
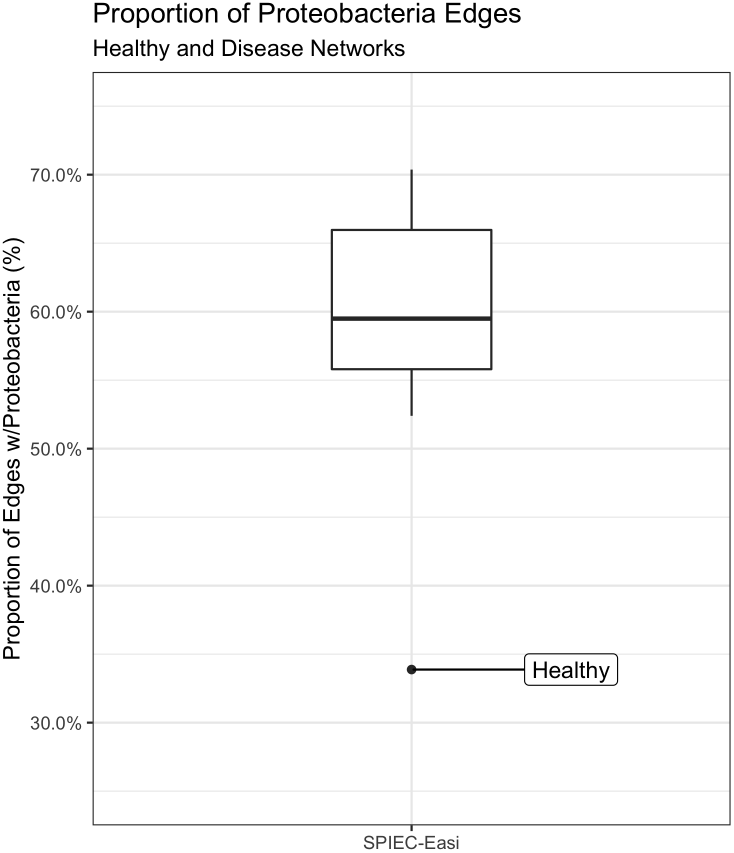
Distribution of the fractions of interactions that involve Proteobacteria in healthy and diseased microbiome association networks. In the boxplot, the Y-axis represents the proportion of interactions involving Proteobacteria.

Taking a closer look at Proteobacteria edges within our networks, we found that nondisease Proteobacteria interactions were often connecting modules pre-dominantly interconnected with Proteobacteria containing edges that were also found in the healthy network (Figure 6). This result shows that beyond microbial co-occurrences commonly shared between healthy and diseased networks, the diseased networks also contain disease-only edges that greatly interconnect Proteobacteria species compared to the healthy network. Additionally, majority of Proteobacteria containing edges within the consensus network were filtered out, being observed in less than 5 networks, suggesting that many of these Proteobacteria connections are not universal across all diseases. Together, this may suggest that the enrichment of Proteobacteria edges observed in disease networks are contributed by rare disease-specific edges, and provide greater interconnectivity between Proteobacteria containing edges that would be otherwise be considered loosely connected when compared to the healthy network.

**Figure 6:**
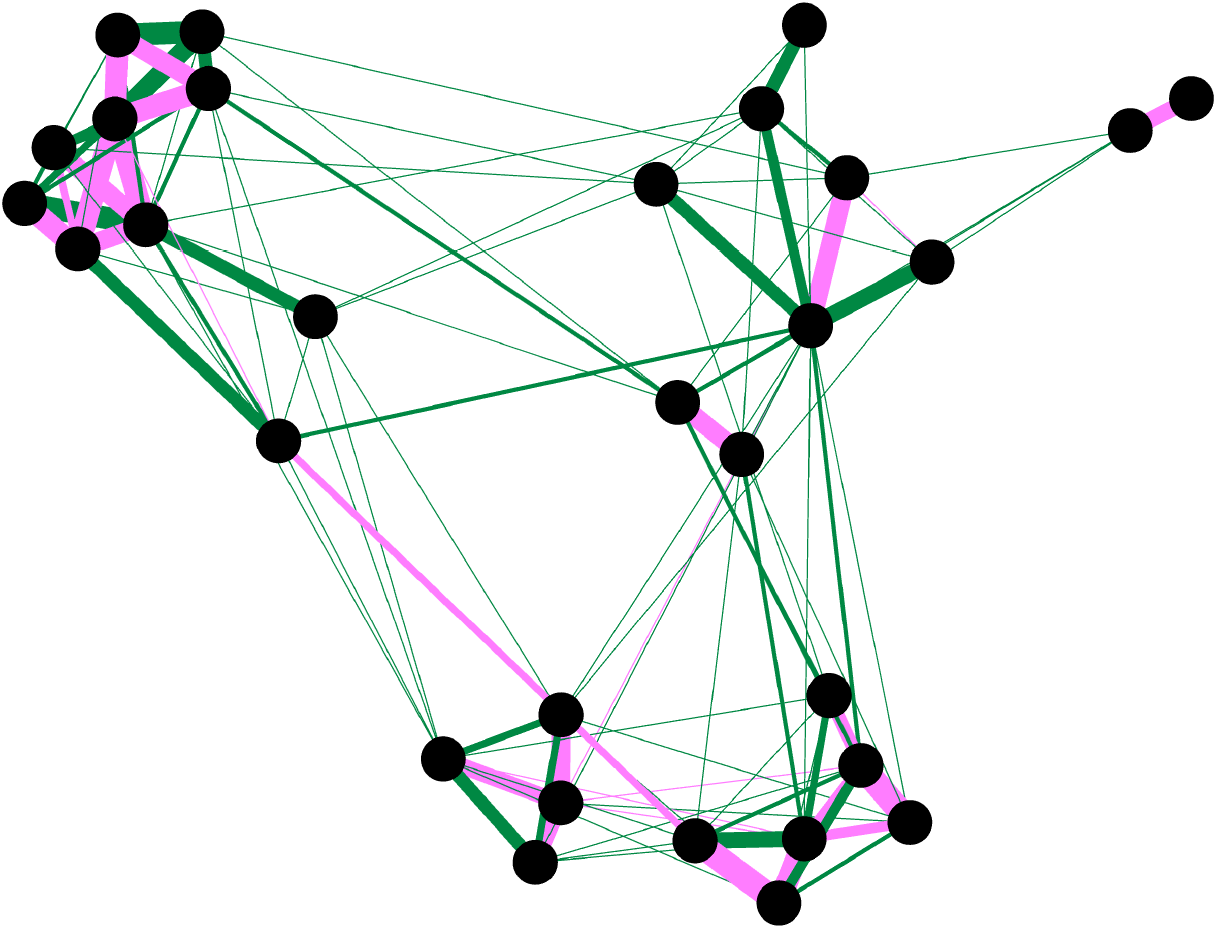
Subgraph containing the largest Proteobacteria module found in consensus network. Consensus network contains edges shared between 5 or more disease networks. Green edges represent edges that are not found in the healthy network, while pink edges represent edges in the consensus network that are also found in the healthy network. Edge weight is scaled by the count of networks that a given edge is observed in.

## Discussion

Here in this meta-analysis of gut microbiome datasets, we report patterns of microbiome interaction within and between healthy and diseased microbiomes through the use of microbiome association networks. Our analysis showed that rare taxa of microbiome datasets can play a disproportionate role in microbiome interactions relative to their taxonomic abundances. While Bacteroidetes and Firmicutes were found to comprise a majority of the microbiome abundances in all microbiome phenotypes, the proportion in which Bacteroidetes and Firmicutes participated in significant network associations in terms of both nodes and edges were unproportional to their high relative abundances. Instead, majority of the significant edges within the microbiome association networks were composed of rarer taxa. This contrast supports previous studies that suggests that rare species may play an overproportional role in microbiome community dynamics compared to their more abundant counterparts[51, 52, 53].

In our observations, we also found several notable differences between healthy and diseased microbiome networks. These observations include an enrichment of Bacteroidetes-Bacteroidetes and Firmicutes-Firmicutes interactions within the Healthy Network and enriched Proteobacteria-Actinobacteria interactions in Diseased Networks. While it is unclear if these differences in association are causal or a result of a diseased state, these differences in interactions highlight dysbiosis in diseased microbiome association networks and can be used as potential markers. Additionally, Diseased network edges were found to be highly enriched for Proteobacteria compared to the Healthy network. The Healthy network had a significantly lower proportion of Proteobacteria participation in association networks compared to Diseased networks, and suggests that increased Proteobacteria interactions with other members of the microbiome may be a hallmark of microbiome dysbiosis. Many of the features identified in this study that stratify between healthy and disease networks were found to be consistent across disease networks, suggesting that these features are not disease-specific but general markers of dysbiosis and features of diseased gut microbiota.

Additionally, by identifying community modules within both Healthy and Diseased networks, we were able to identify community modules that were resilient to change and the community interactions that were likely to be retained across different microbiome association networks regardless of phenotypic association. While these modules do not necessarily represent a ‘core’ microbiome associated with a particular phenotype, these resilient modules help us better understand the underlying microbiome community structure that is shared between phenotypes. Investigation into better understanding of microbiome community structure and assembly dynamics can help future efforts in modulating the human gut microbiome. Module resilience highlights the advantages of meta-analysis, and utilizing standardized approaches so that cross-disease and cross-study analysis can be generalized across datasets to help us better understand microbiome dynamics spanning across diseased states.

While this analysis did not include all possible studies or diseases, this study highlights the benefits of re-analyzing studies with standardized procedures so that results can be generalizable and compared between datasets. That being said, there still remains much limitations to microbiome meta-analyses and microbiome interaction as a whole. In particular, as there often lacks widely accepted reference standard and adopted protocol, methods and techniques utilized to analyze microbiome data is widely left open to interpretation and researchers can only inform themselves of the nuances between methods and select the method that best fits their data, needs, and available resources. Issues of possible variation and confounding factors such as experimental or sequencing artifacts, environmental factors, batch effect, differences in taxonomic annotation and quantification methods, technical artifacts highly limit robust downstream analysis. To mitigate some of the potential issues of confounding factors (e.g. species-level annotation error, batch effect between studies, variation between study design and patient selection), we focused majority of our analysis by agglomerating at the Phylum level, but acknowledge there remains much more to be explore at lower taxonomic levels. There remains an increasing need for gold standards to be developed so that tools and methods can be benchmarked and evaluated to establish standardized protocols. Future efforts in development of experimental and computational methods are necessary to address issues of microbiome compositionality.

Despite these limitations, our results uncovered features of microbiome association between healthy and diseased cohorts that may help future efforts in understanding alterations of the gut microbiome that may be associated with diseased states. Here we provide all microbial association networks produced as part of this study as a resource for future efforts in studying microbial associations. While it is not possible to assess and benchmark the wide availability of microbial association methods, standardizing the protocols and processing steps of data analysis will help future efforts to uncover features that warrant further investigation. By preforming meta-analyses, results of individual studies can reach beyond itself and assist in contextualizing new results through expanding insights in comparison to other studies. Nevertheless, computational microbiome association methods remain insufficient by themselves to identify causal interactions. Association analysis can only serve as a starting point to reduce the search space and identify potential candidates for downstream hypothesis testing and experimental validation.

## Data Availability

All code, metadata, and graph files generated as part of this study is available in the GutNet repository located at https://github.com/mgtools/GutNet.

## Acknowledgements

This work was supported by NIH grant 1R01AI143254 and NSF grant 2025451. The authors thank Dr. Yong Yeol Ahn for helpful discussions of network based analyses.

## Author Contributions

T.L. and Y.Y. conceived the research. T.L and Y.Y wrote programs to process data. T.L. processed the data and preformed all analysis. T.L. and Y.Y. interpreted results and participated in preparing the manuscript.

