## Supplementary Figures for "Meta-analysis of Microbiome Association Networks Reveal Patterns of Dysbiosis in Diseased Microbiomes"

### Supplementary File

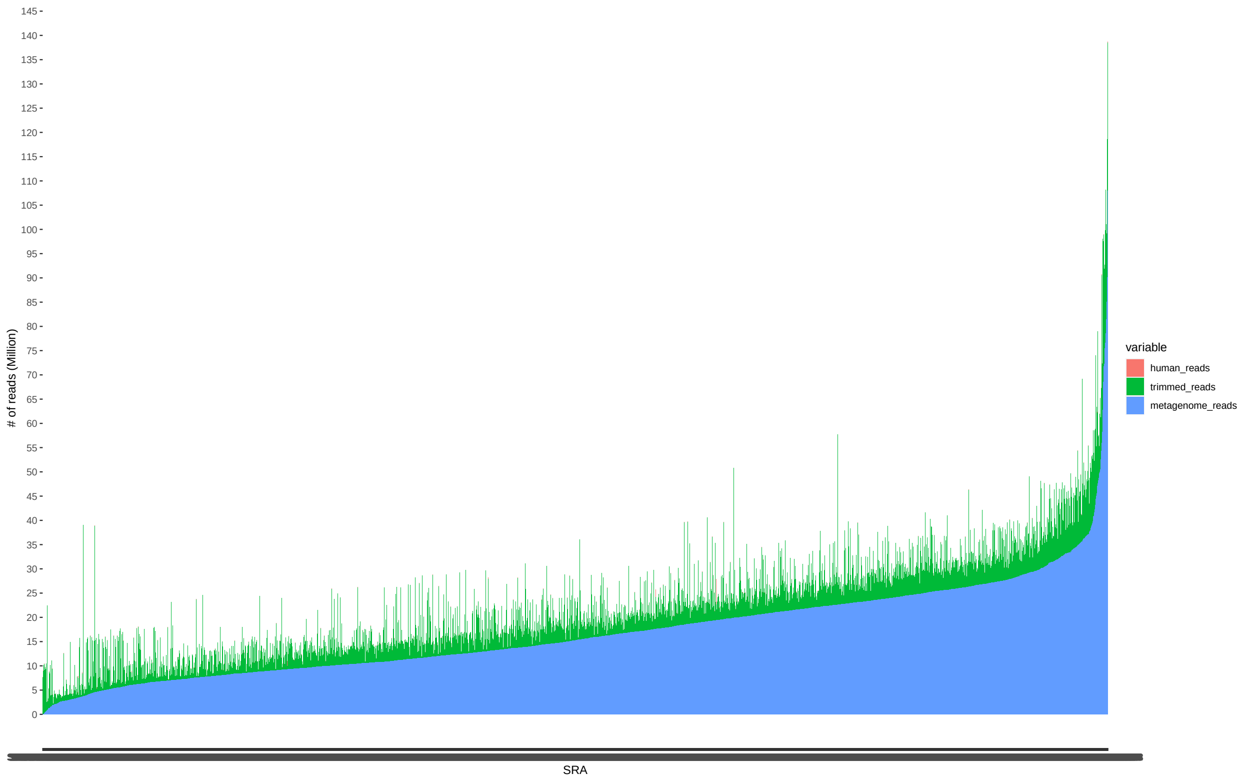


Supplementary Figure 1: Distribution of reads per sample. Blue reads include reads maintained after quality trimming and host filtering. Green reads represent proportion of reads that were quality trimmed. Red reads represent reads that were contaminated human reads. X-axis represents individual SRA included in this analysis. Y-axis represent number of reads per sample, scaled to reads-per-million (RPM).

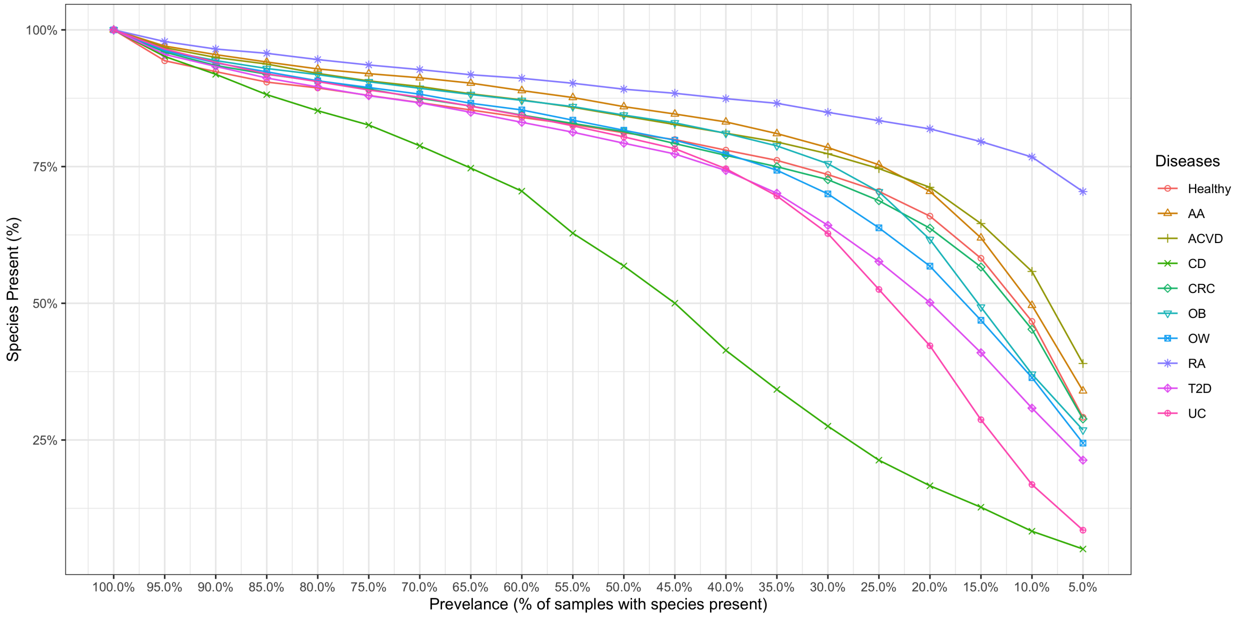


Supplementary Figure 2: Identified species retained decreases with increasing prevalence threshold. The X-axis represents the prevalence threshold to filter species at in increments of 5%. The Y-axis represents the proportion of species represented in a given disease phenotype abundance matrix.

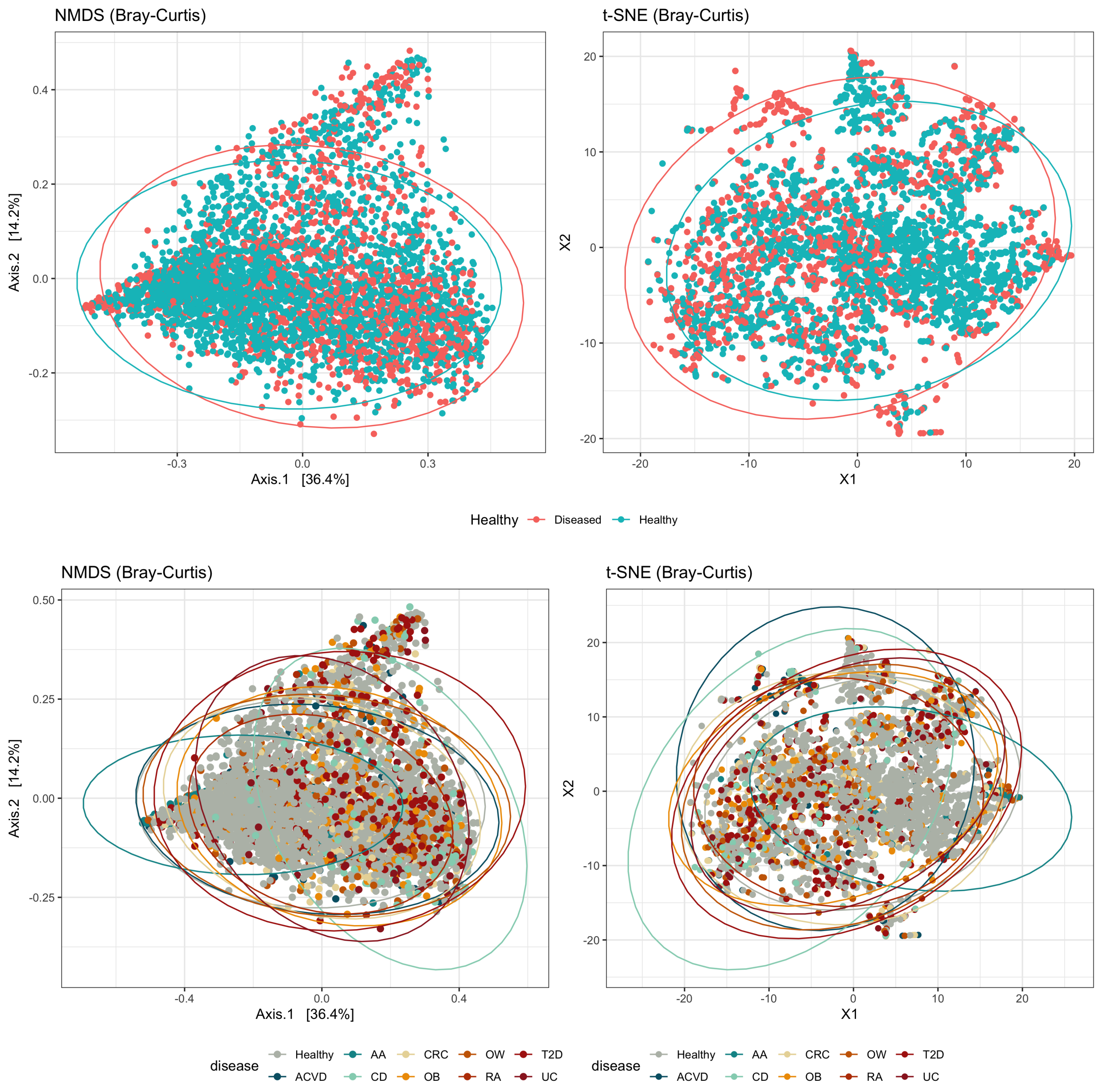


Supplementary Figure 3: Two-dimensional projections of the microbiome samples based on their species profiles. (A) non-metric multidimensional scaling (NMDS) plots based on Bray–Curtis dissimilarity between species profiles colored by healthy and diseased samples, (B) t-SNE plot based on Bray-Curtis dissimilarity measure colored by healthy and diseased samples, (C) NMDS plots based on Bray-Curtis dissimilarity measure colored by phenotype, and (D) t-SNE plot based on Bray-Curtis dissimilarity measure colored by phenotype.

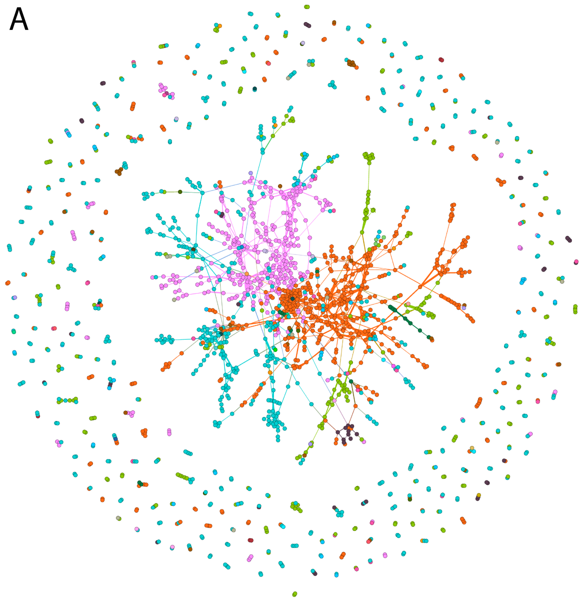


Supplementary Figure 4: Microbiome association networks of healthy samples constructed using SPIEC-EASI. (A) nodes colored by Phylum-level taxonomic assignment, (B) nodes colored by community modules predicted by Leiden algorithm.

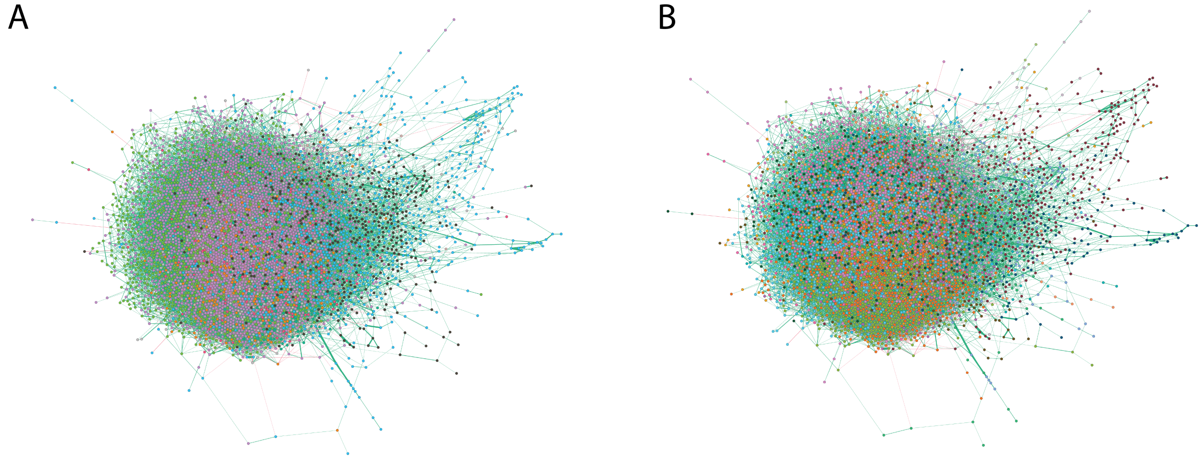


Supplementary Figure 5: Advanced Adenoma (AA) association network. (A) Network are colored by a node's taxonomic annotation at the Phylum level. (B) Network are colored by their community partition predicted by the Leiden algorithm. Edge weights are assigned by their association value inferred by SPIEC-EASI, where green edges represent positive edges, and negative edges represent negative edges.

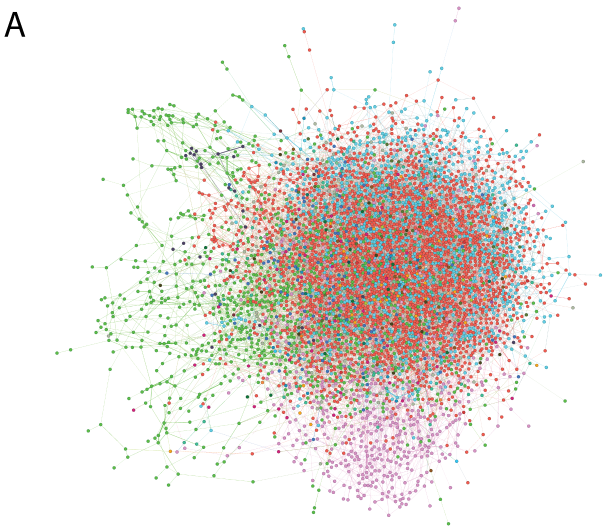


Supplementary Figure 6: Atherosclerotic Cardiovascular Disease (ACVD) association network. (A) Network are colored by a node's taxonomic annotation at the Phylum level. (B) Network are colored by their community partition predicted by the Leiden algorithm. Edge weights are assigned by their association value inferred by SPIEC-EASI, where green edges represent positive edges, and negative edges represent negative edges.

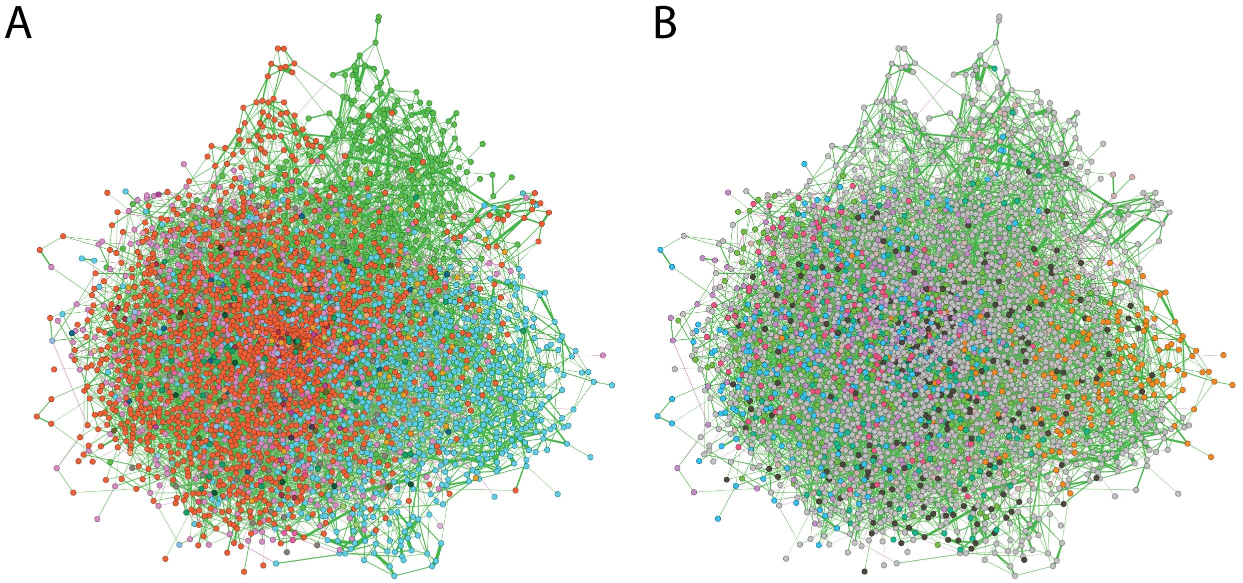


Supplementary Figure 7: Colorectal Cancer (CRC) association network. (A) Network are colored by a node's taxonomic annotation at the Phylum level. (B) Network are colored by their community partition predicted by the Leiden algorithm. Edge weights are assigned by their association value inferred by SPIEC-EASI, where green edges represent positive edges, and negative edges represent negative edges.

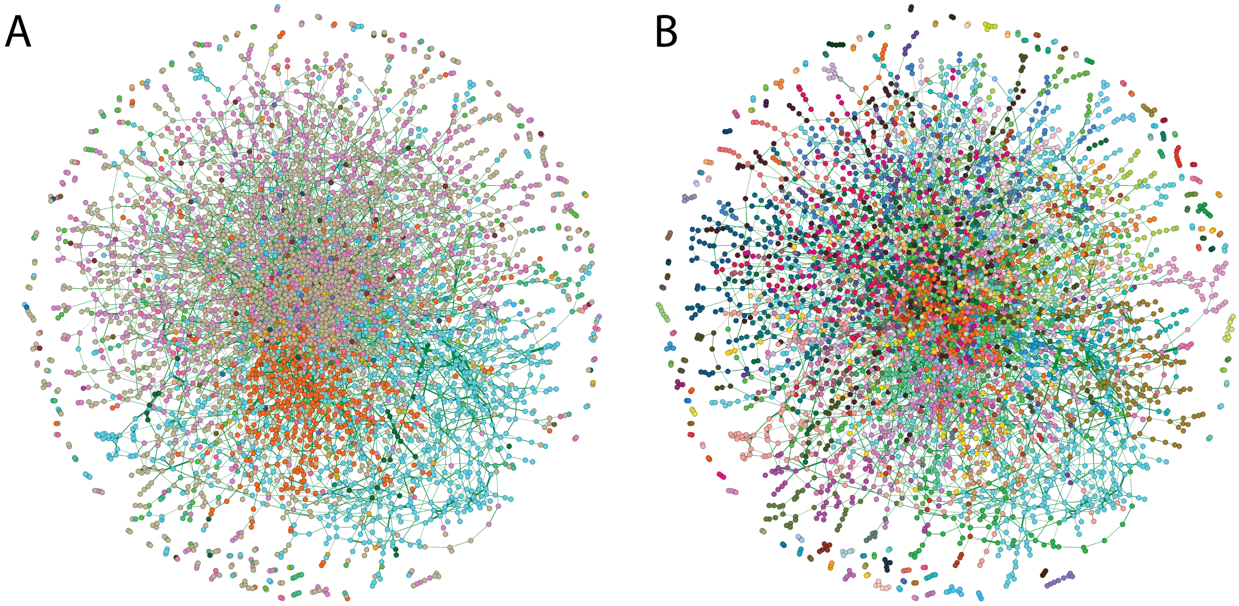


Supplementary Figure 8: Crohn's Disease (CD) association network. (A) Network are colored by a node's taxonomic annotation at the Phylum level. (B) Network are colored by their community partition predicted by the Leiden algorithm. Edge weights are assigned by their association value inferred by SPIEC-EASI, where green edges represent positive edges, and negative edges represent negative edges.

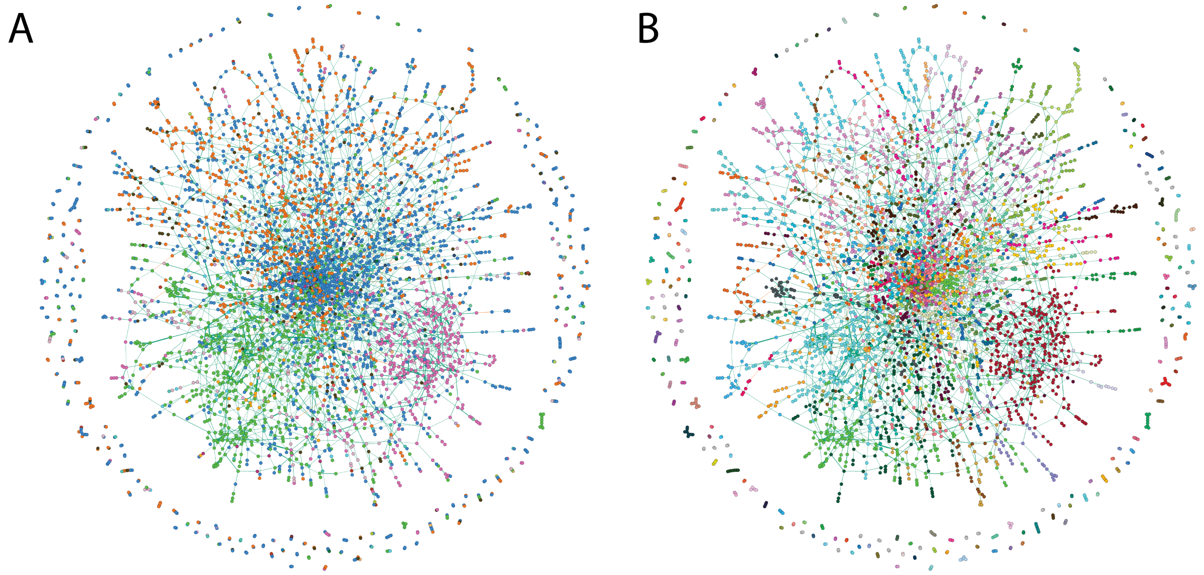


Supplementary Figure 9: Obesity (OB) association network. (A) Network are colored by a node's taxonomic annotation at the Phylum level. (B) Network are colored by their community partition predicted by the Leiden algorithm. Edge weights are assigned by their association value inferred by SPIEC-EASI, where green edges represent positive edges, and negative edges represent negative edges.

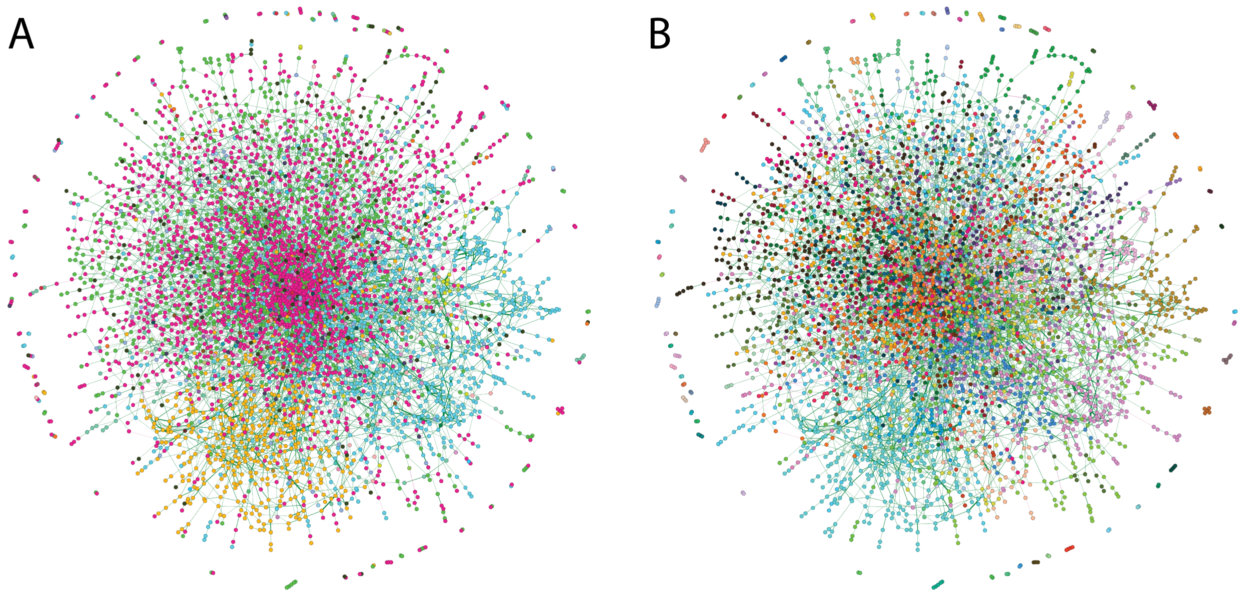


Supplementary Figure 10: Overweight (OW) association network. (A) Network are colored by a node's taxonomic annotation at the Phylum level. (B) Network are colored by their community partition predicted by the Leiden algorithm. Edge weights are assigned by their association value inferred by SPIEC-EASI, where green edges represent positive edges, and negative edges represent negative edges.

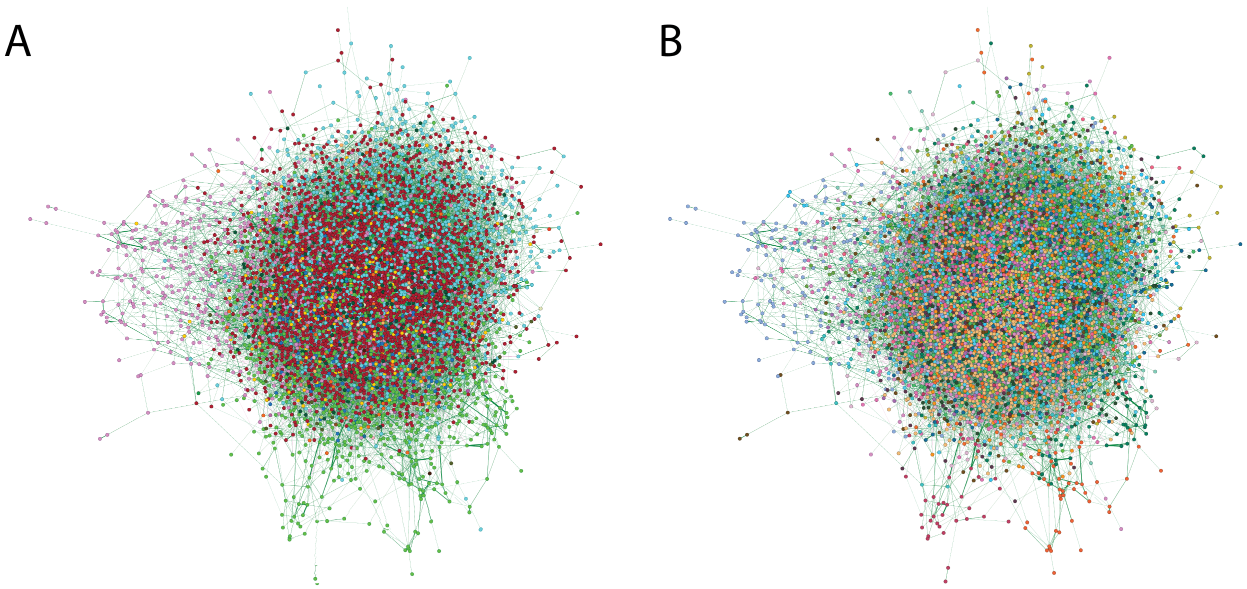


Supplementary Figure 11: Rheumatoid Athritis (RA) association network. (A) Network are colored by a node's taxonomic annotation at the Phylum level. (B) Network are colored by their community partition predicted by the Leiden algorithm. Edge weights are assigned by their association value inferred by SPIEC-EASI, where green edges represent positive edges, and negative edges represent negative edges.

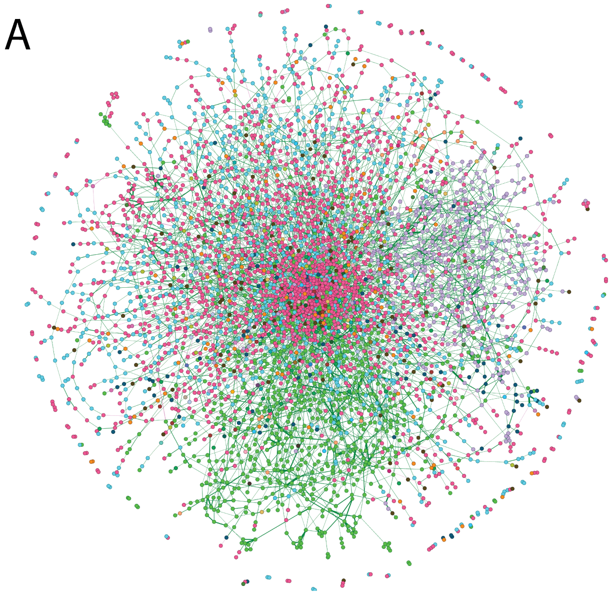


Supplementary Figure 12: Type 2 Diabetes (T2D) association network. (A) Network are colored by a node's taxonomic annotation at the Phylum level. (B) Network are colored by their community partition predicted by the Leiden algorithm. Edge weights are assigned by their association value inferred by SPIEC-EASI, where green edges represent positive edges, and negative edges represent negative edges.

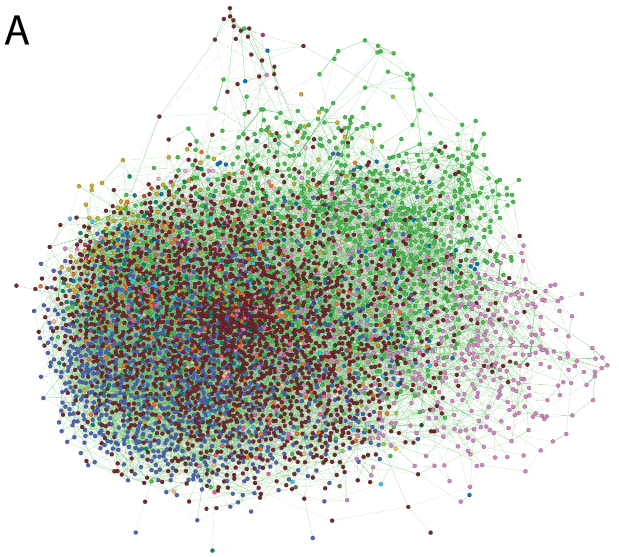


Supplementary Figure 13: Ulcerative Colitis (UC) association network. (A) Network are colored by a node's taxonomic annotation at the Phylum level. (B) Network are colored by their community partition predicted by the Leiden algorithm. Edge weights are assigned by their association value inferred by SPIEC-EASI, where green edges represent positive edges, and negative edges represent negative edges.

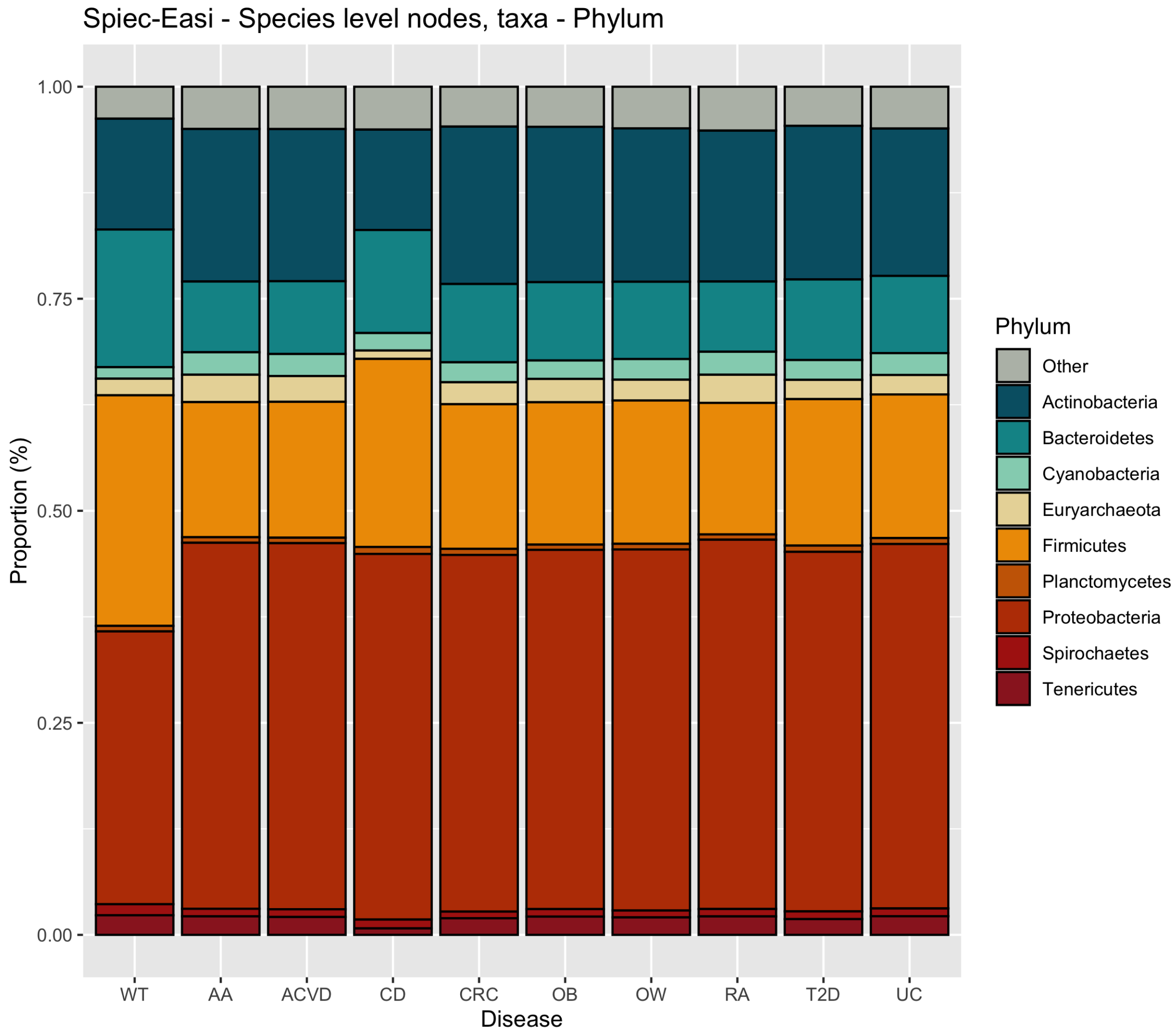


Supplementary Figure 14: Distribution of node by taxonomic assignment per each phenotypic network.
